# Spectrin Interactome under normal and HbE-disease conditions

**DOI:** 10.1101/2020.10.03.324822

**Authors:** Dipayan Bose, Sk Ramiz Islam, Sutapa Saha, Abhijit Chakrabarti

## Abstract

Spectrin, the major component of the erythrocyte membrane skeleton is a key player in red cell biology. It has a significant role in signalling pathways and as such knowledge of spectrin interactors becomes crucial. Here we report the cytosolic interactome of human erythroid spectrin (ProteomeXchange id: PXD021525). This is to the best of our knowledge the first report of the interactome of human erythroid spectrin. We have further investigated the spectrin interactome under HbE disease conditions. Our findings indicate that there is no difference in the identity of the proteins interacting with spectrin between normal and disease conditions. However relative abundance of the interacting partners is seen to change. Very interestingly the interacting chaperone proteins, heme-containing proteins and redox active proteins are seen to be up-regulated in HbE disease state. This is consistent with our previous observation that presence of oxidation prone hemoglobin variants leads to an increase of redox regulators and chaperones in the red cell proteome. Spectrin can also interact with horse radish peroxidase and oxidatively crosslink hemoglobin, which has possible implications in oxidative stress management. Since a large fraction of spectrin interacting proteins are chaperones and redox active proteins, it is possible that spectrin may have a broader role in redox regulation, especially in cases where there are unstable hemoglobin variants present.

## INTRODUCTION

Spectrin is the major component of the membrane skeleton in mature human erythrocytes (RBCs) (Bennett 1985). It is known that spectrin has a role as a component of the cellular signalling machinery, and is also involved as a structural platform for cytoskeletal protein assemblies (Djinovic-Carugo, Gautel et al. 2002, Deng, Wang et al. 2015, Fletcher, Elbediwy et al. 2015).

Moreover studies from our lab have demonstrated that spectrin is able to interact with hemoglobin, the most abundant protein of the RBC cytosol (Basu and Chakrabarti 2015). Such hemoglobin interactions seem to prefer structurally perturbed hemoglobin variants like hemoglobin E (HbE) over normal adult hemoglobin A (HbA) (Datta, Chakrabarty et al. 2003).

Spectrin HbE interactions are redox active and it is found that the two proteins can oxidatively crosslink in the presence of hydrogen peroxide (Datta, Basu et al. 2006). This is important as spectrin is shown to be able to act as a chaperone for hemoglobin (Basu and Chakrabarti 2015).

Indeed the chaperone like activity of spectrin extends to other non related redox active proteins such as horse radish peroxidase, and other proteins such as insulin and BSA (Chakrabarti, Bhattacharya et al. 2001, Bose, Patra et al. 2017). Interestingly, the chaperone activity of spectrin is found to favour hemoglobin over other proteins (Bose and Chakrabarti 2019).

Spectrin is also known to interact with proteins such as actin and ankyrin via specific sites on its extended rod-like surface (Bennett 1989, Ipsaro, Huang et al. 2009). Spectrin surface is also implicated in its interaction with hemoglobin, where a “beads on a string” model is proposed (Mishra, Chakrabarti et al. 2016).

Further, defects in spectrin are implicated in the physiology of RBC diseases like hereditary spherocytosis and elliptocytosis (Liu, Derick et al. 1990). Such diseases are thought to be a manifestation of deficits in the protein-protein interaction capacity of spectrin.

These protein-protein interaction driven biological roles of spectrin indicate the importance of an understanding of the interacting partners of spectrin. Given the affinity of spectrin for HbE over HbA, and its possible role in redox biology, we hypothesize that the spectrin interactome may change in E-disease conditions (majority of hemoglobin is of the HbE variant,) where the cytosolic proteome is known to differ in amounts of chaperones and redox regulators (Bhattacharya, Saha et al. 2010). In this study we have presented a report of the cytosolic interacting partners of spectrin, both in normal conditions and E-disease states.

## MATERIALS AND METHODS

Normal adult human blood samples containing hemoglobin variant HbA were collected from healthy volunteers with proper informed consent. Blood samples from homozygous HbE patients were obtained from Ramkrishna Mission Seva Pratisthan Hospital, Kolkata, India, with informed consent of the patients following the guidelines of the Institutional Ethical Committee. Samples were characterized using Bio-Rad Variant HPLC system and variant hemoglobin content was determined. Only those samples with HbE content greater than 95% were used. For HbA and HbE, a total of 5 samples each, were pooled together to reduce biological variability.

### Hemoglobin depletion

Hemoglobin depleted blood lysate was prepared by the method of Ringrose *et. al.* (Ringrose, van Solinge et al. 2008). Human RBCs were lysed via incubation against 20 volumes of hypotonic buffer (5 mM Na-phosphate, pH 8.0) for 12 hours at 4°C, and resulting lysate was cleared of membrane debris via centrifugation. 200 mg of the resulting soluble cytosolic protein fraction was taken in 2 ml of buffer containing 50 mM phosphate, pH 8.0, 300 mM NaCl, and 5 mM imidazole. It was run on 8 ml Ni-NTA Super flow resin at 0.2 ml/min at 4 °C. The flow-through containing Hb depleted cytosolic proteins were pooled and concentrated.

### Spectrin preparation and immobilization

Human erythrocytic spectrin was prepared as described earlier (Basu and Chakrabarti 2015). Purified spectrin was attached to CNBr activated Sepharose resin using the protocol of Kavran *et. al.* (Kavran and Leahy 2014).

Briefly, human erythroid spectrin was dialyzed into cold coupling buffer (NaHCO3, 100 mM, pH 8.3, NaCl 500 mM) at 4 °C, and then concentrated to 2 mg/ml. For each 1 mg of dialyzed spectrin, 0.25 mg of dry CNBr resin was hydrated by addition of 5 column-volumes of cold activation buffer (1 mM HCl) on a rocker at 4 °C for 2 hours. Swollen resin was centrifuged at 1000g for 5 minutes and supernatant was discarded. Spectrin in coupling buffer was added and incubated overnight at 4 °C on a rocker. Resin was then centrifuged and supernatant collected to calculate fraction of bound spectrin. Reaction was quenched with quenching buffer (100 mM Tris-HCl, pH 8.0) and column was washed and equilibrated with binding buffer (50 mM Na-phosphate buffer pH 7.4, 100 mM NaCl). It was calculated that 1mg protein was bound per ml of resin. Some spectrin free resin was quenched in quenching buffer for later control steps.

### Pull down assay

10 mg of hemoglobin depleted cytosolic protein fraction was taken in 5 ml of binding buffer (50 mM Na-phosphate buffer pH 7.4, 100 mM NaCl) and incubated with 1 ml of spectrin bound CNBr Sepharose overnight at 4 °C. Prey bound resin was washed extensively with 20 column volumes of the same buffer to remove non-specifically bound proteins. Prey proteins were eluted in 20% acetic acid, 1 % SDS.

As control, 10 mg of depleted cytosolic fraction was run on spectrin free quenched CNBr resin similarly and after washing was eluted with 20% acetic acid, 1% SDS. The final eluate was collected and analysed on 12% SDS gel along with those of the test samples.

For each of HbA and HbE samples, eluates of three distinct experiments were pooled together, concentrated and dialysed against 100 mM Tris-HCL pH 8.0 and 6M GdmCl and supplied as such to Valerian Chem. Pvt. Ltd., New Delhi, India, who analysed our samples.

### Mass Spectrometric analysis

25 μl samples were taken and reduced with 5 mM TCEP and further alkylated with 50 mM iodoacetamide and then digested with trypsin (1:50, Trypsin/lysate ratio) for 16 h at 37 °C. Digests were cleaned using a C18 silica cartridge to remove the salt and dried using a speed vac. The dried pellets were resuspended in buffer A (5% acetonitrile, 0.1% formic acid).

For both the HbA and HbE spectrin interactome pools, LC-MS/MS was run were performed thrice to reduce variability among runs. All the experiments were performed using EASY-nLC 1000 system (Thermo Fisher Scientific) coupled to Thermo Fisher-QExactive equipped with nanoelectrospray ion source. 1.0 μg of the peptide mixture was resolved using 25 cm PicoFrit column (360μm outer diameter, 75μm inner diameter, 10μm tip) filled with 1.8 μm of C18-resin (Dr Maeisch, Germany). The peptides were loaded with buffer A and eluted with a 0–40% gradient of buffer B (95% acetonitrile, 0.1% formic acid) at a flow rate of 300 nl/min for 100 min. MS data was acquired using a data-dependent top10 method dynamically choosing the most abundant precursor ions from the survey scan.

All samples were processed and RAW files generated were analyzed with Proteome Discoverer (v2.2) against the Uniprot Human reference proteome database. For Sequest search, the precursor and fragment mass tolerances were set at 10 ppm and 0.5 Da, respectively. The protease used to generate peptides, i.e. enzyme specificity was set for trypsin/P (cleavage at the C terminus of “K/R: unless followed by “P”) along with maximum missed cleavages value of two. Carbamidomethyl on cysteine was set as fixed modification and oxidation of methionine and N-terminal acetylation and phosphorylation on site (S, T, Y) were considered as variable modifications for database search. Both peptide spectrum match and protein false discovery rate were set to 0.01 FDR. The mass spectrometry proteomics data have been deposited to the ProteomeXchange Consortium via the PRIDE (Perez-Riverol, Csordas et al. 2019) partner repository with the dataset identifier PXD021525.

The quantitative interactome analysis involved label-free relative protein quantification, following LC-MS, using the Minora Feature Detector Node of the Proteome Discoverer 2.2 with default settings. The peptide spectrum matches with high confidence were only considered. Unique and Razor peptides were used for label-free quantification. Peptide precursor abundance was determined using peptide intensity. Normalization was automatically applied by Proteome Discoverer software using total peptide amount option. Imputation was not selected during Proteome Discoverer Data Processing pipeline. A criterion of a minimum of two peptides per protein was not applied to data analyzed by Orbitrap technology.

## RESULTS AND DISCUSSIONS

### Pull down assay

The analysis of the pull down fractions on 12% SDS gel showed that Ni-NTA resin depleted the vast majority of Hb and spectrin free quenched CNBr resin did not bind RBC cytosolic proteins. Further, spectrin interacting proteins generated distinctly different patterns on the gel than Hb depleted cytosolic fraction. A representative picture is shown in **Figure 1**.

**Figure 1:**
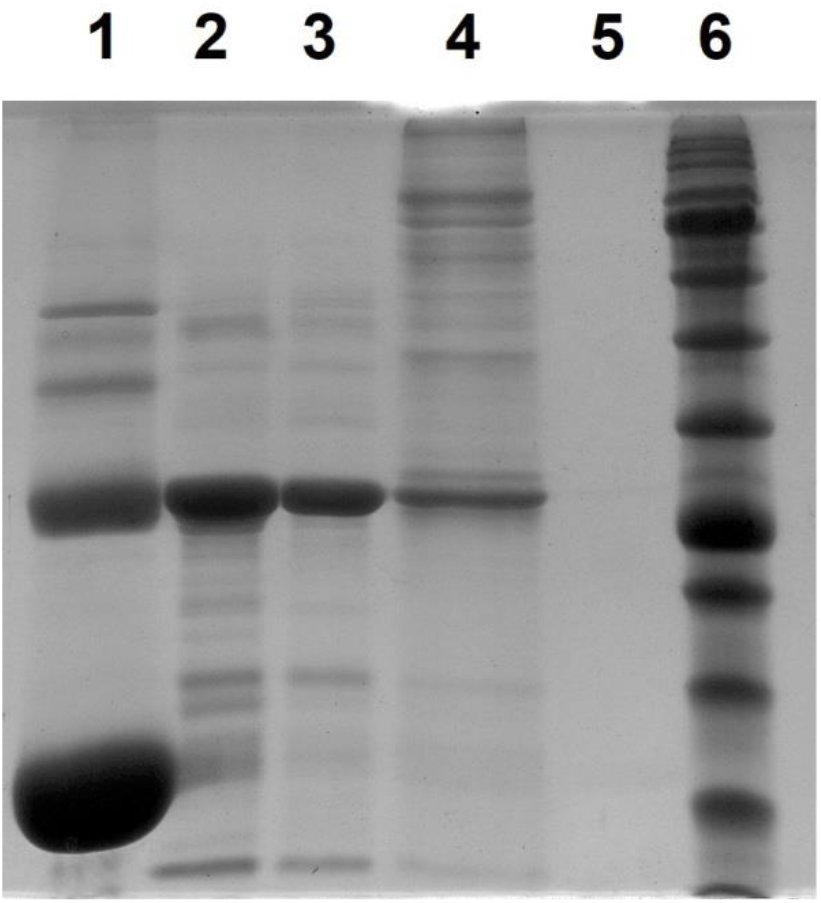
Figure shows silver stained 12% SDS gel of the different RBC cytosolic fractions used for interactome analysis. Lane 1 shows crude RBC lysate, bottom most band is for hemoglobin, the next most prominent band is for carbonic anhydrase. Lane 2 shows cytosolic fraction after hemoglobin depletion using Ni-NTA. Lane 3 shows the flow through washings of the prey pool after binding CNBr Sepharose immobilized spectrin. Lane 4 shows the spectrin interacting proteins eluted with 20% acetic acid, 1% SDS. Lane 5 shows the same elution with prey pool incubated with control CNBr quenched with Tris-HCl. No detectable proteins were found to bind the resin free of spectrin. Lane 6 shows marker (Prism Ultra, Abcam).

### Mass Spectrometric Analysis

The generated list of spectrin interactors was validated against a quantitative list of complete RBC proteome by Bryk *et. al.* (Bryk and Wiśniewski 2017). Contaminant proteins, keratin and HSA were ignored, as was hemoglobin variants. Only proteins with 2 or more unique peptides, high FDR score, Sequest hit score greater than 100 and CV% less than 20 were considered. In this way 214 proteins were identified. Data was independently validated using Mascot software. List of proteins is given in **Supplementary Table S1**.

The final protein list after shorting was also used to generate a volcano plot and heat-map in the MetaboAnalyst (Chong, Wishart et al. 2019) online statistical server. The data were sum-normalized, log-transformed, and Pareto-scaled within the server.

For the volcano-plot, a fold change threshold and FDR adjusted P-value was set to 2 and 0.05 respectively to find significant changes in protein abundance between the groups. The plot is shown in **Supplementary Figure S1.**

It is important to note that there were no differences in the number or identity of proteins interacting with spectrin in both cases; however the relative abundance of the interactors had changed. Spectrin was found to significantly interact with heme-containing redox proteins, chaperones, lipid modulating proteins, membrane skeletal proteins, protein quality control maintenance machinery and signaling components. **Figure 2** shows a pie chart of the different protein classes the interactome of spectrin can be divided into.

**Figure 2:**
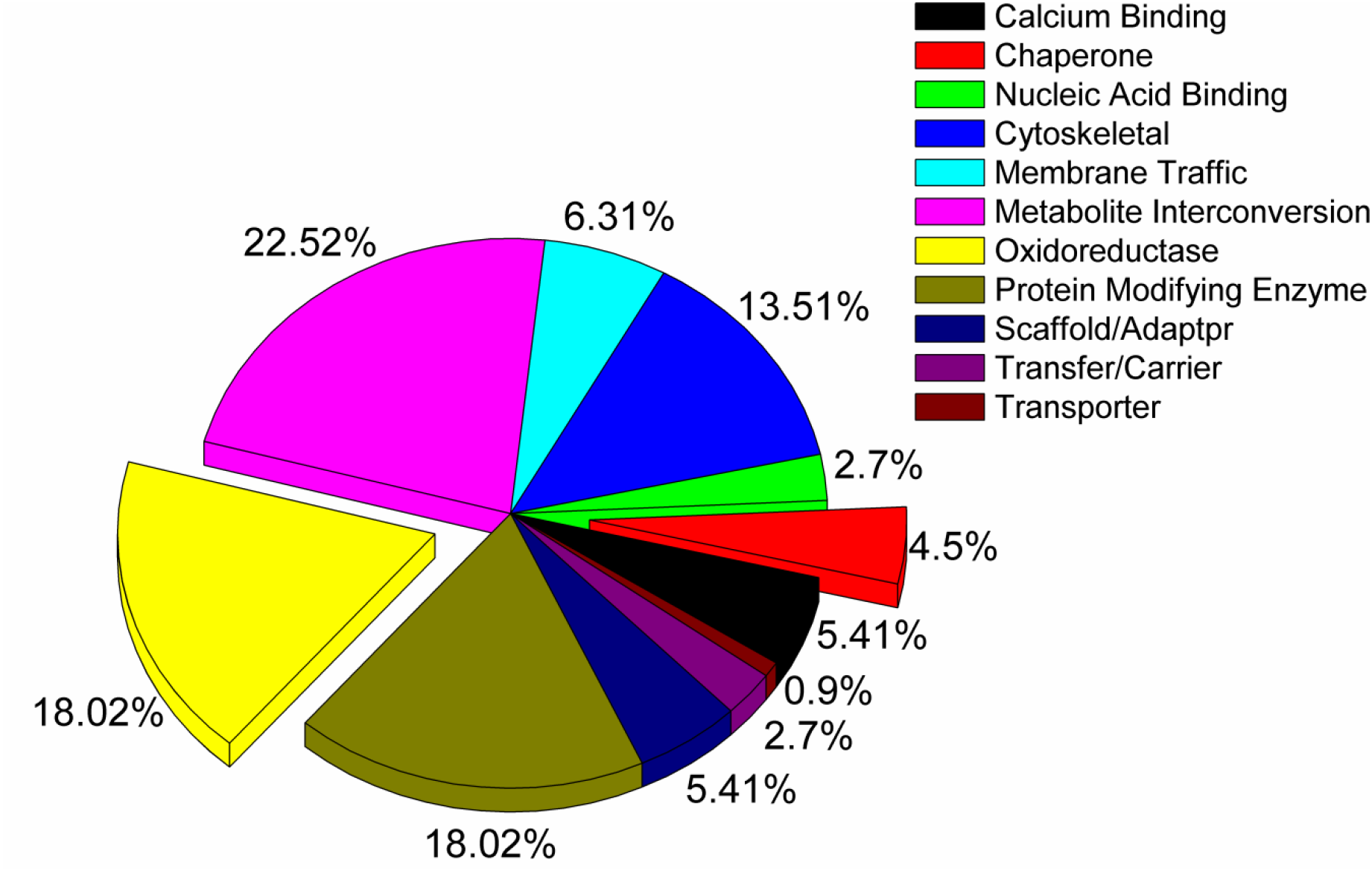
Pie chart representing the different classes of proteins that spectrin interacting proteins belong to. Classification of proteins into different classes was done with the help of PANTHER database.

A major class of spectrin interactor is seen to be redox active proteins, many of which are also heme containing. Interestingly these proteins are differentially up or down-regulated in hemoglobin E-disease than in normal states. This is in line with our previous observations where we showed Eβ-thalassemia caused an up-regulation of cytosolic redox regulators and chaperones (Chakrabarti, Halder et al. 2016) caused due to unstable hemoglobin variants mediated oxidative stress leading to accumulation of anti-oxidant machinery (Basu, Saha et al. 2013). **Table 1** tabulates the representative set of spectrin interacting redox active proteins and chaperones, and a heat-map of these representative proteins showing altered abundance between the groups is presented in **Figure 3**. Red color indicates the highest relative concentration and green color indicates lower relative concentration.

**Figure 3:**
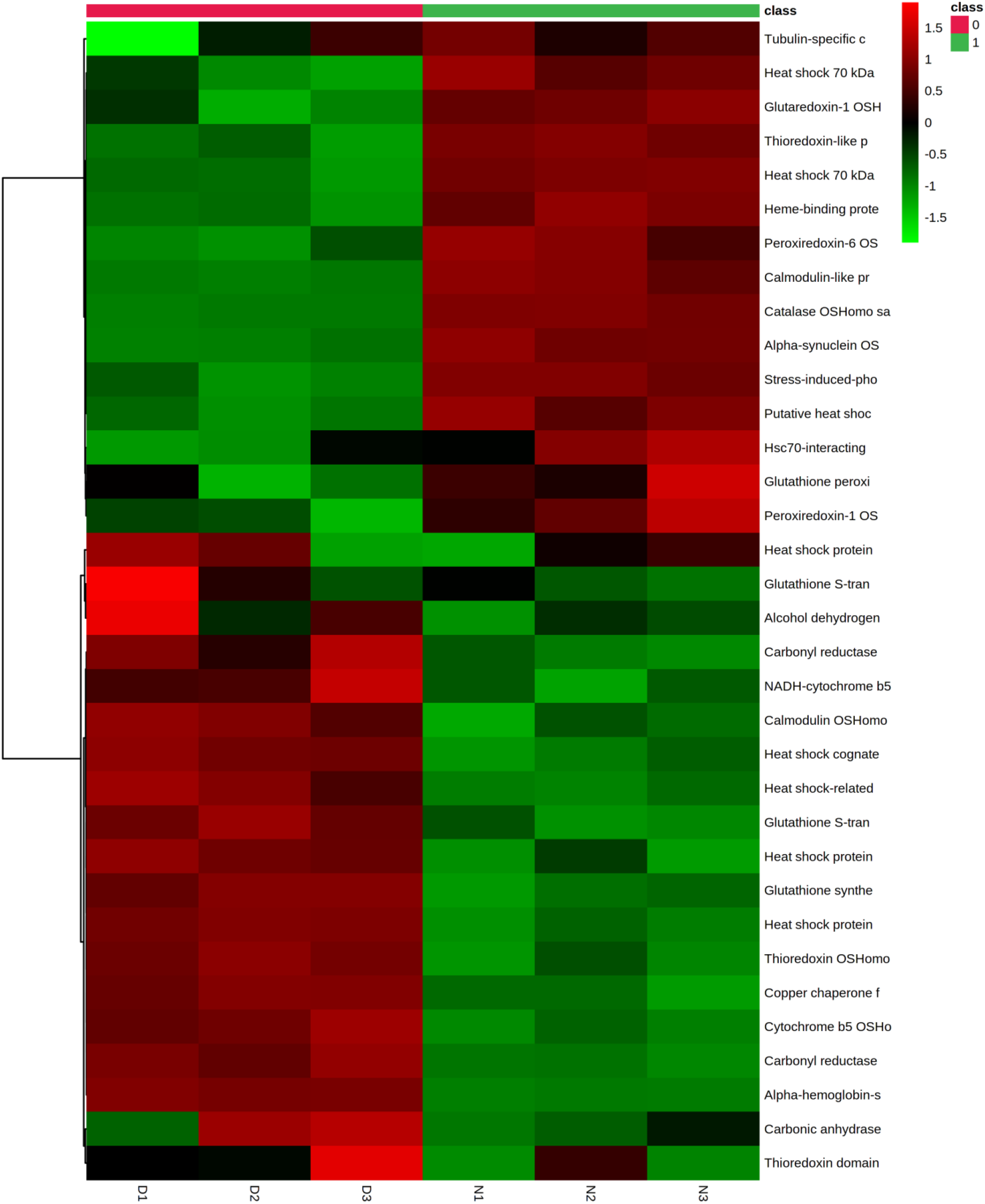
A heat map of the proteins tabulated in **Table 1** is shown. The samples numbered D1, 2 and 3 are the three individual LC-MS/MS runs of the HbE-disease state samples and those numbered N1, 2 and 3 are those for normal HbA state samples. It is seen that there is differential up regulation of some redox active and chaperone proteins in the disease state over that of normal conditions.

**Table 1:**
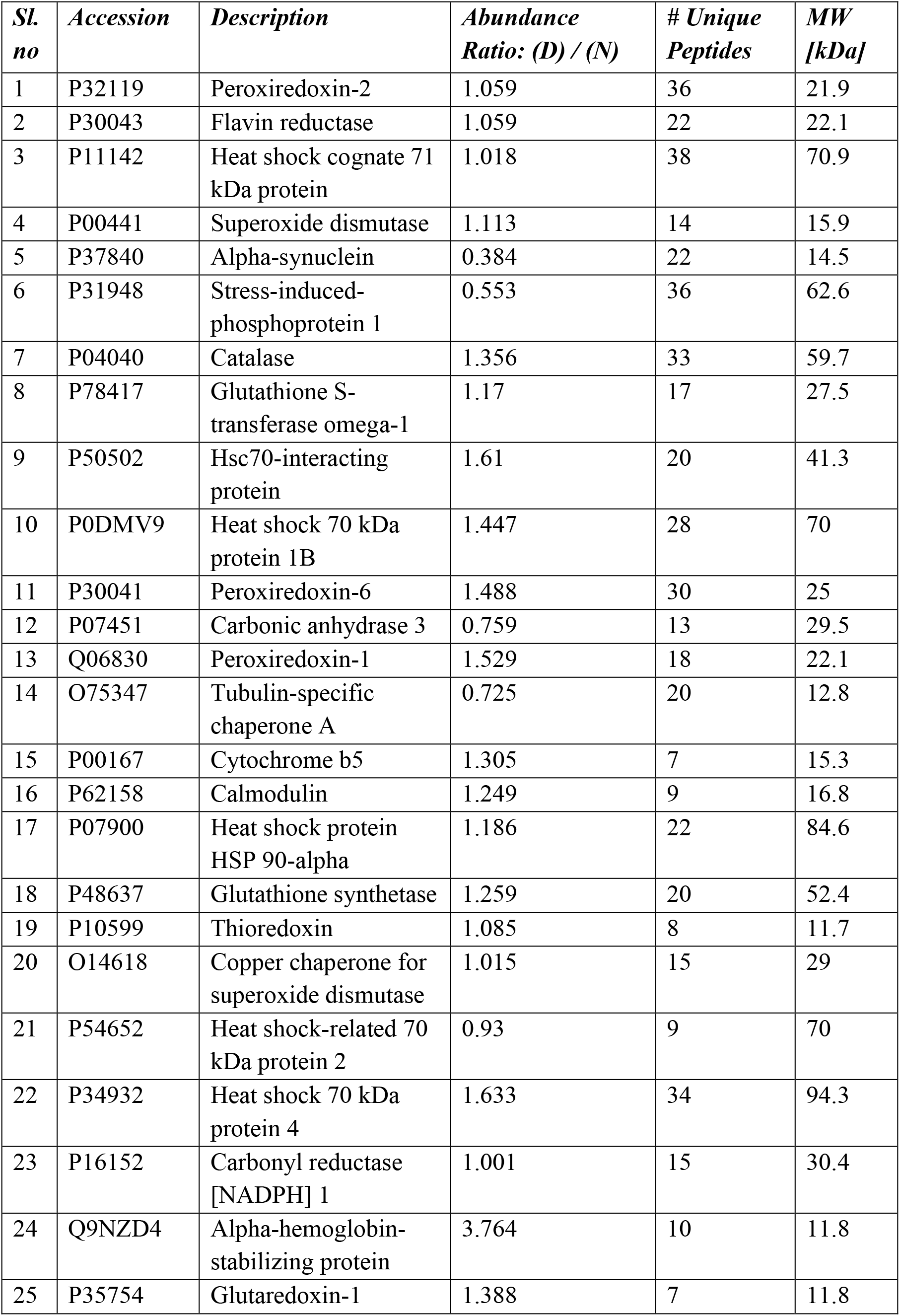
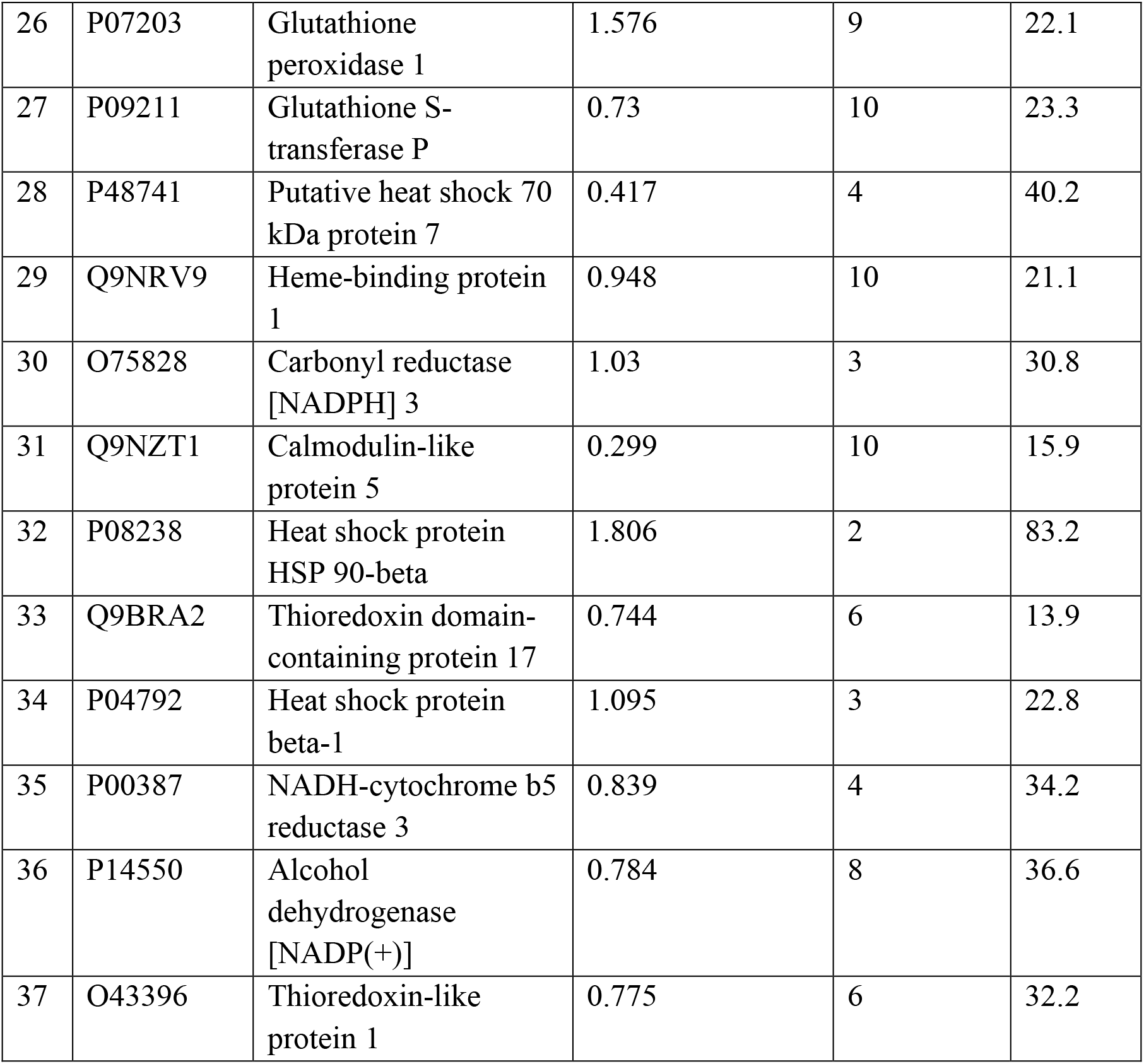
A representative set of the redox active proteins and chaperone proteins found interacting with spectrin are tabulated. Their relative abundances in disease state - hemoglobin E-disease (D) versus their relative abundance in normal state – containing hemoglobin A (N) is given as D/N.

| *Sl. no* | *Accession* | *Description* | *Abundance Ratio: (D) / (N)* | *# Unique Peptides* | *MW [kDa]* |
| --- | --- | --- | --- | --- | --- |
| 1 | P32119 | Peroxiredoxin-2 | 1.059 | 36 | 21.9 |
| 2 | P30043 | Flavin reductase | 1.059 | 22 | 22.1 |
| 3 | P11142 | Heat shock cognate 71 kDa protein | 1.018 | 38 | 70.9 |
| 4 | P00441 | Superoxide dismutase | 1.113 | 14 | 15.9 |
| 5 | P37840 | Alpha-synuclein | 0.384 | 22 | 14.5 |
| 6 | P31948 | Stress-induced-phosphoprotein 1 | 0.553 | 36 | 62.6 |
| 7 | P04040 | Catalase | 1.356 | 33 | 59.7 |
| 8 | P78417 | Glutathione S-transferase omega-1 | 1.17 | 17 | 27.5 |
| 9 | P50502 | Hsc70-interacting protein | 1.61 | 20 | 41.3 |
| 10 | P0DMV9 | Heat shock 70 kDa protein 1B | 1.447 | 28 | 70 |
| 11 | P30041 | Peroxiredoxin-6 | 1.488 | 30 | 25 |
| 12 | P07451 | Carbonic anhydrase 3 | 0.759 | 13 | 29.5 |
| 13 | Q06830 | Peroxiredoxin-1 | 1.529 | 18 | 22.1 |
| 14 | O75347 | Tubulin-specific chaperone A | 0.725 | 20 | 12.8 |
| 15 | P00167 | Cytochrome b5 | 1.305 | 7 | 15.3 |
| 16 | P62158 | Calmodulin | 1.249 | 9 | 16.8 |
| 17 | P07900 | Heat shock protein HSP 90-alpha | 1.186 | 22 | 84.6 |
| 18 | P48637 | Glutathione synthetase | 1.259 | 20 | 52.4 |
| 19 | P10599 | Thioredoxin | 1.085 | 8 | 11.7 |
| 20 | O14618 | Copper chaperone for superoxide dismutase | 1.015 | 15 | 29 |
| 21 | P54652 | Heat shock-related 70 kDa protein 2 | 0.93 | 9 | 70 |
| 22 | P34932 | Heat shock 70 kDa protein 4 | 1.633 | 34 | 94.3 |
| 23 | P16152 | Carbonyl reductase [NADPH] 1 | 1.001 | 15 | 30.4 |
| 24 | Q9NZD4 | Alpha-hemoglobin-stabilizing protein | 3.764 | 10 | 11.8 |
| 25 | P35754 | Glutaredoxin-1 | 1.388 | 7 | 11.8 |
| 26 | P07203 | Glutathione peroxidase 1 | 1.576 | 9 | 22.1 |
| 27 | P09211 | Glutathione S-transferase P | 0.73 | 10 | 23.3 |
| 28 | P48741 | Putative heat shock 70 kDa protein 7 | 0.417 | 4 | 40.2 |
| 29 | Q9NRV9 | Heme-binding protein 1 | 0.948 | 10 | 21.1 |
| 30 | O75828 | Carbonyl reductase [NADPH] 3 | 1.03 | 3 | 30.8 |
| 31 | Q9NZT1 | Calmodulin-like protein 5 | 0.299 | 10 | 15.9 |
| 32 | P08238 | Heat shock protein HSP 90-beta | 1.806 | 2 | 83.2 |
| 33 | Q9BRA2 | Thioredoxin domain-containing protein 17 | 0.744 | 6 | 13.9 |
| 34 | P04792 | Heat shock protein beta-1 | 1.095 | 3 | 22.8 |
| 35 | P00387 | NADH-cytochrome b5 reductase 3 | 0.839 | 4 | 34.2 |
| 36 | P14550 | Alcohol dehydrogenase [NADP(+)] | 0.784 | 8 | 36.6 |
| 37 | O43396 | Thioredoxin-like protein 1 | 0.775 | 6 | 32.2 |

For network analysis, String (v 11.0) was used. The network was built using a high confidence interaction score (0.700). A total number of 203 nodes were found with 882 edges. The PPI enrichment P-value was found to be less than 1.0e-16, which indicates the number of interactions is more than random interaction with a random set of proteins of similar size. Such enrichment signifies that the proteins are at least partially involved in a particular set of biological functions. The network is shown in **Supplementary Figure S2.**

## CONCLUSIONS

Spectrin is known to interact with hemoglobin and heme-containing HRP and act as their chaperone (Bhattacharyya, Ray et al. 2004). Recently we have generated preliminary data that shows spectrin interaction causes enhancement of the peroxidase activity of heme proteins (manuscript under review); this has implications in the clearance of ROS and oxidative stress. The observation that spectrin also interacts with several other heme-containing proteins and proteins involved in heme management, as well as redox regulatory proteins lends support to the idea that spectrin may be involved in oxidative stress management in RBCs.

## ABBREVIATIONS

HbA: Hemoglobin A
HbE: Hemoglobin E

## Ethics Approval

Institutional Ethical Committee of Ramkrishna Mission Seva Pratisthan Hospital, provided ethical approval.

## Data Availability

Datasets for MS experiments are available on PRIDE under identifier PXD021525.

## Code Availability

N/A

### Author’s Contributions

All authors contributed significantly to the writing of manuscript and performance of experiments/data acquisition.

