## Supplementary figures for "Spectrin Interactome under normal and HbE-disease conditions"

**Figure S1:**

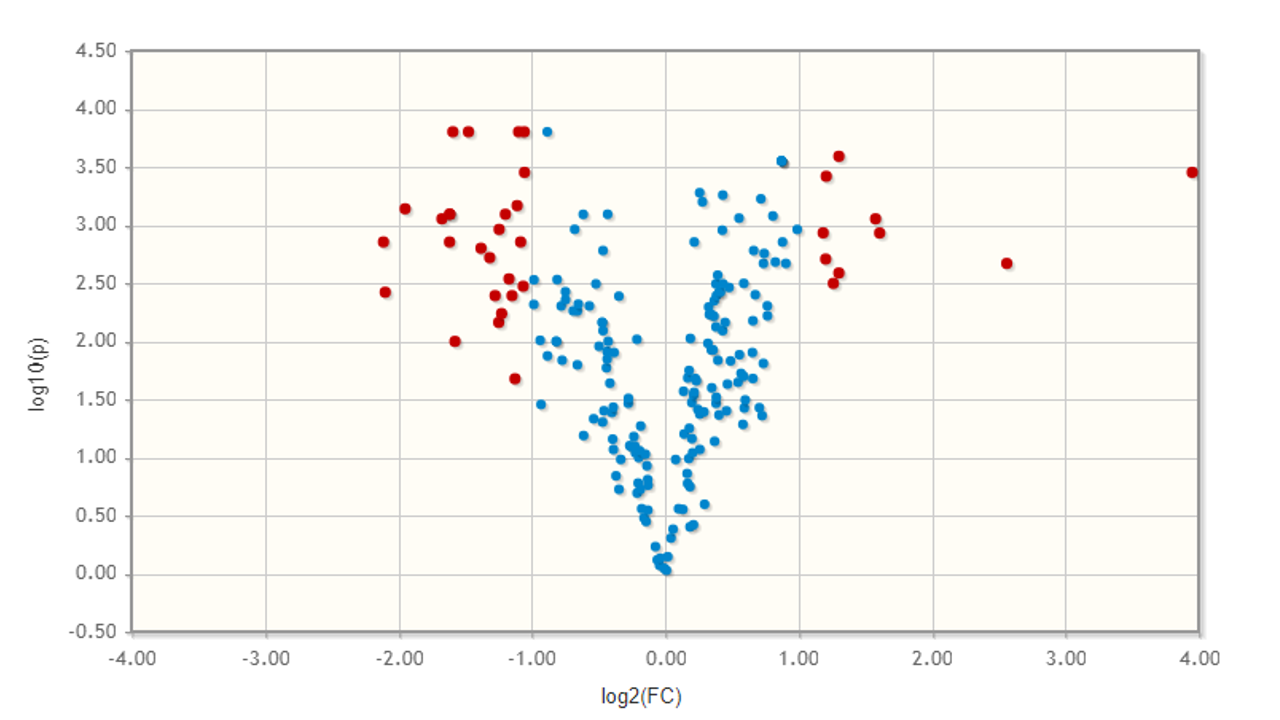

**Figure S1:** Volcano plot shows significant deviation in the relative abundance of proteins between HbA and HbE samples.

**Figure S2:**

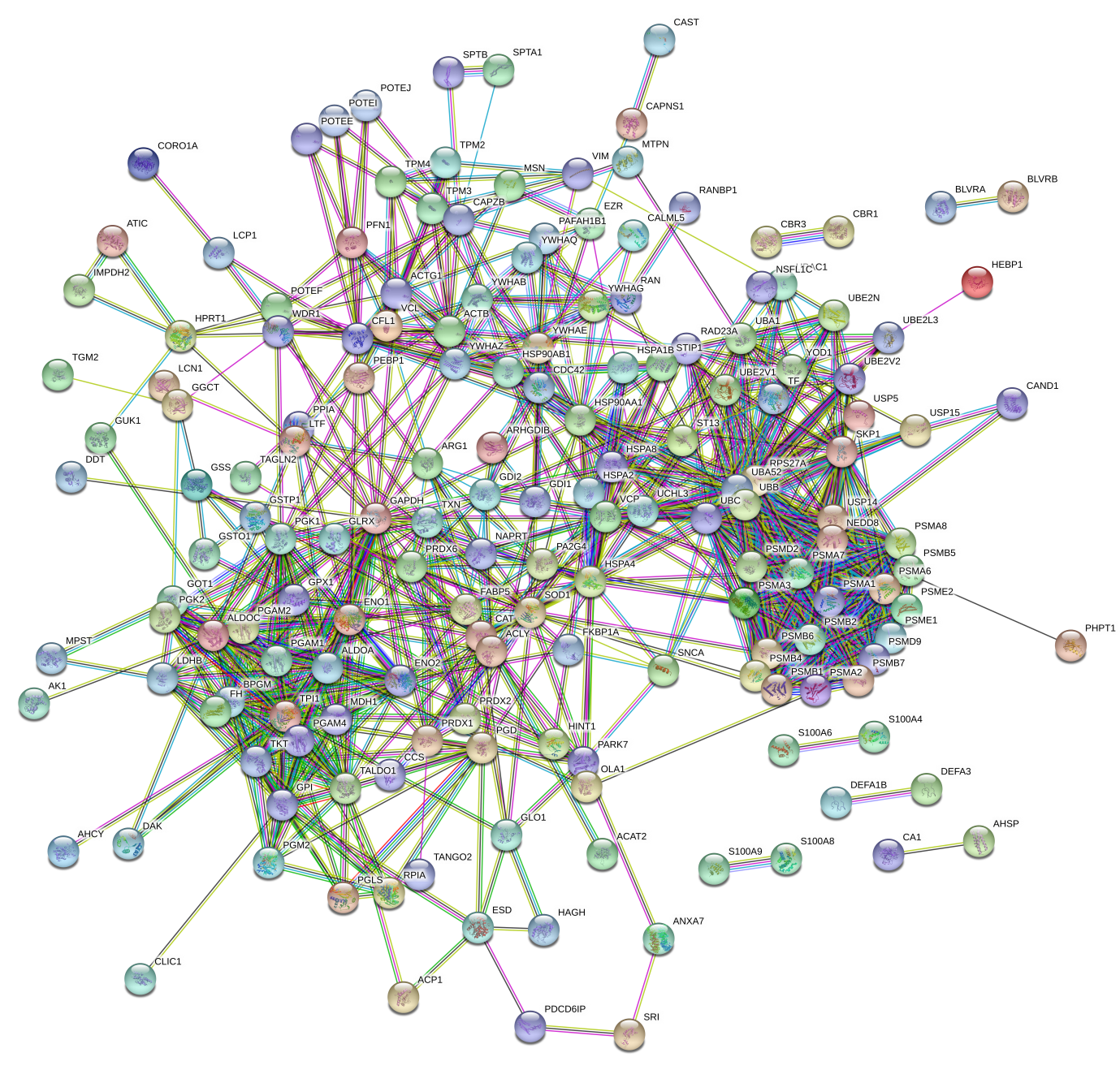

**Figure S2:** String (v 11.0) was used to create a network of spectrin interacting proteins using a high confidence interaction score (0.700). A total number of 203 nodes were found with 882 edges. The PPI enrichment P-value was found to be less than 1.0e-16, which indicates the number of interactions is more than random interaction with a random set of proteins of similar size.
